# High relatedness in sexually produced restoration cohorts of the endangered elkhorn coral, *Acropora palmata*

**DOI:** 10.64898/2026.07.15.738610

**Authors:** Carly D. Kenkel, Holland Elder, Grace McDermott, Trinity Conn, Nicolas S. Locatelli, Iliana B. Baums, Courtney Klepac, Zachary Craig, Dakotah Merck, R. Scott Winters, Margaret W. Miller, Dana E. Williams, Erinn M. Muller

## Abstract

Climate change is driving the decline of coral populations around the world such that many are unlikely to recover without human intervention. Assisted sexual reproduction is one intervention proposed to enhance genetic diversity and support population recovery, yet its genetic outcomes remain poorly quantified. We evaluated genome-wide relatedness and genetic diversity in 168 restoration genets of the endangered Caribbean coral *Acropora palmata*, including 153 sexually produced offspring derived from multi-parent batch and biparental crosses. We detected high relatedness within multi-parent batch cross cohorts, with many genets comprising only one or a few full-sibling groups, indicating highly unequal parental contributions. Nucleotide diversity was lower in one batch cross cohort relative to founder populations, but the absolute difference was small and runs of homozygosity were relatively short indicating that inbreeding depression is not yet a concern. These patterns suggest that common larval propagation approaches can successfully generate large numbers of new genets but underscore the need to manage inbreeding risk, especially in small breeding stocks such as the Caribbean *Acropora* spp. Specifically, our results highlight the need for comprehensive genetic management to integrate assisted sexual reproduction into coral restoration, including parentage tracking, broodstock rotation, and relatedness-informed outplanting designs.

## Introduction

Climate change is accelerating the global loss of species (Urban 2015; Ceballos et al. 2015) with increasing temperatures as a major driver of decline (Román-Palacios and Wiens 2020). A recent example of this is the fourth global coral bleaching event, which resulted in significant coral population declines globally due to record high ocean temperatures in 2023-25 (Reimer et al. 2024; Neely et al. 2024; Ceccarelli et al. 2026). For populations already at risk, such as the critically endangered elkhorn coral, *Acropora palmata* (Williams et al. 2008, 2020), the results were dire. In 2025, *A. palmata* was declared functionally extinct in Florida (Manzello et al. 2025). To prevent total extirpation of this economically and ecologically important species, there is a recognized need for population management interventions to halt and/or reverse both the demographic and genetic declines (Rodriguez-Clark et al. 2025).

Population management aims include increasing numbers of individuals for reintroduction into the wild and/or maintaining genetic diversity (Mace and Ballou 1990). Such programs have been successfully applied to halt declines in captive zoo and aquarium populations (Che-Castaldo et al. 2021) and even restore wild populations once considered extinct in the wild, as in the famous case of the black-footed ferret (Livieri et al. 2022). Equal representation of founders (i.e., the initial genetically unique individuals originating from the natural population) is generally a top priority to minimize both random genetic drift and domestication selection leading to adaptation to the artificial conditions of *ex situ* facilities (Rodriguez-Clark et al. 2025). For species which experience many sexually reproduced generations in human care, this involves carefully monitored pedigrees, breeding plans and reintroduction designs to limit the potential for inbreeding depression due to matings among close relatives.

Methodological advances in coral mariculture (Craggs et al. 2017; O’Neil et al. 2021; Koch et al. 2022) have now made it possible to integrate assisted breeding into coral conservation and restoration programs (Randall et al. 2020; Vardi et al. 2021; Banaszak et al. 2023) (Fig. 1A). However, most population management plans were designed for vertebrates, which produce far fewer offspring per breeding pair than mass-spawning marine invertebrates, like corals, capable of producing billions of gametes annually (Harrison and Wallace 1990; Harrison 2011) which can yield hundreds to thousands of offspring per cross. In addition, historical effective population sizes for most coral species, including Caribbean acroporids, were likely quite large (Prada et al. 2016; Cramer et al. 2020; Fuller et al. 2020; Cooke et al. 2020). Large, outbred populations harbour larger numbers of recessive deleterious genetic variants, making them more susceptible to genetic risks arising from inbreeding following population contraction (Charlesworth and Willis 2009). Consequently, while the high fecundity of coral can accelerate demographic recovery by generating many new individuals, there is a correspondingly elevated risk of inbreeding depression arising from the potential for high relatedness within these sexually produced cohorts.

**Figure 1.**
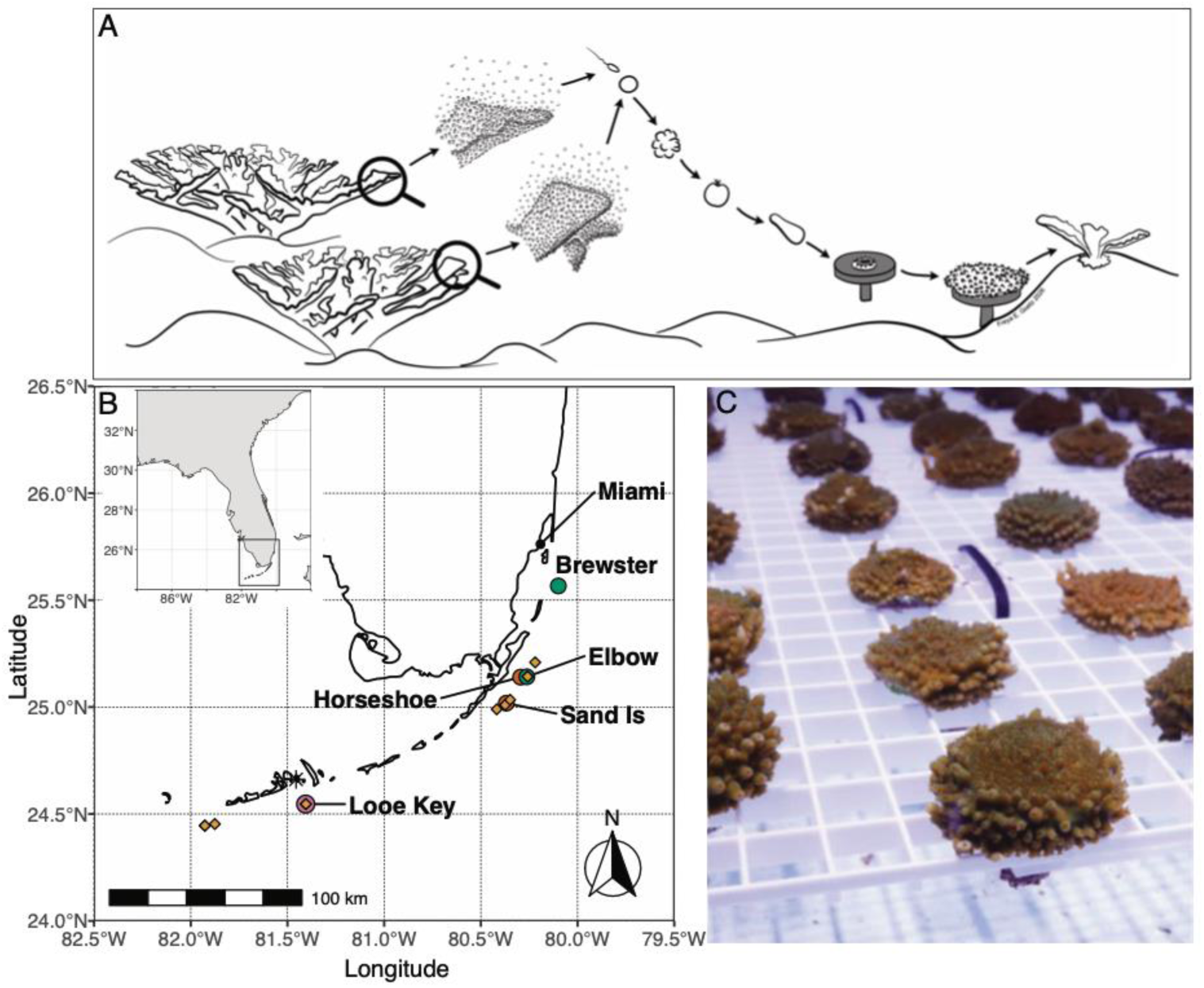
Florida restoration populations of *Acropora palmata*. (A) Reproductive biology of *Acropora palmata* and application to sexual reproduction for restoration. Gametes are collected during annual spawning events and combined to yield larvae, which are settled onto individual plugs and clonally propagated. Illustration by Freya E. Goetz. (B) Map of source populations. Circles indicate reef site origin of parents used in batch (Elbow-Brewster/Ball Buoy and Elbow-Horseshoe-Sand Island) and biparental (Looe Key) crosses. Diamonds indicate coordinates of additional founder genets sampled for high coverage whole genome sequencing and the asterisk indicates the location of the IC2R3 *ex situ* coral nursery. (C) Example photograph of common gardened *A. palmata* ramets in the IC2R3 *ex situ* nursery.

Methods for generating sexually produced cohorts for coral restoration include targeted two parent or biparental crosses (Fig. 1A), where offspring are expected to be full siblings, or multi-parent ‘batch’ crosses in which gametes from several founders are mixed yielding variable levels of relatedness among offspring (Baums et al. 2013, 2022). Although greater genetic diversity may be expected in offspring of multi-parent batch crosses, Baums et al. (2022) report high relatedness in a small outplanted population of sexually produced *A. palmata* genets in Curacao, with the majority of outplants comprising full and half-siblings. Similarly, in an experimental lab-bred population of *A. hyacinthus*, López-Nadam et al. (López-Nandam et al. 2022) also report outsize contributions of specific parents to the resulting offspring cohort, with the majority of parents leaving no surviving offspring. In restoration programs where the parentage of offspring is not actively monitored, the silent nature of parental dropout may mask losses in genetic diversity from restored populations. Consequently, there is a need to determine the extent of these genetic bottlenecks in captively-bred coral populations to design improved coral breeding strategies in support of population management planning (Baums et al. 2022).

In support of this need, we used whole-genome resequencing to estimate relatedness and genome-wide genetic diversity in a total of 168 genets of *A. palmata*, of which 153 were sexually produced offspring of wild founders that have been used in reef restoration operations in Florida since 2018 (E. Muller, pers. comm.). Consistent with prior studies, we find that while a large number of offspring genets can be successfully reared from batch crosses for use in reef restoration, these individuals exhibit very high relatedness, meaning that the realized contribution of each parent is lower than the apparent number of individuals included in the cross.

## Materials and Methods

### Coral origin

*A. palmata* coral were sourced from Mote Marine Laboratory’s *ex-situ* coral nursery at the Elizabeth Moore International Center for Coral Reef Research and Restoration (IC2R3) facility in Summerland Key, FL (24.6617° N, -81.4554° W) and the Coral Restoration Foundation’s (CRF) Key Largo nursery (24.98300° N,- 80.435500° W) (Fig. 1B). Mote’s *A. palmata* were composed of 162 genotypes from the coral restoration broodstock. Of these, 153 genotypes were sexually produced offspring of wild founder coral, meaning that they were reared from individual larvae following mass spawning of adult populations and subsequently clonally propagated within *ex-situ* coral nursery aquaria (Fig. 1A,C). The 153 sexually produced genotypes were derived from different mass spawning events (Table S1); specifically, 26 originated from a five-parent batch cross between Elbow, Horseshoe, and Sand Island Reef founder genets in 2013 (hereafter the E-H-SI Batch Cross), 80 from three different batch crosses on sequential nights in 2017, a five-parent batch fertilization of Elbow Reef founders on the night of 10 August, a separate fertilization of coral which spawned at Biscayne National Park (Brewster/Ball Buoy reefs) also on the night of 10 August, and a three-parent cross between founder genets from Elbow Reef the night of 11 August. Larvae from these 2017 fertilizations were subsequently combined in the IC2R3 holding facility and are hereafter referred to as the E-B Batch Cross. An additional 43 sexually produced genets derived from a 2020 biparental cross between two Lower Keys founder genets (hereafter the LK-3 x LK-10 Biparental Cross), and finally, three from a 2014 batch cross comprised solely of Elbow Reef parents and one from a 2015 batch cross among Elbow reef parents (Table S1). The final nine genotypes were asexually propagated from Lower Florida Keys founder genets sourced directly from Looe Key, Sand Key, Western Dry Rocks, and Turtle Reef between 2014-2021 (Fig. 1B). CRF’s *A. palmata* consisted of asexually propagated coral obtained from six founder genotypes sourced from five reefs in the Upper Florida Keys: Carysfort, Elbow, French, Molasses and Pickles (Fig. 1B).

### DNA extraction & sequencing

Holobiont DNA was extracted from snap frozen tissue samples following a modified CTAB-chloroform protocol (Baker and Cunning 2015). Each sample also was cleaned using an isopropanol sodium acetate DNA clean-up to remove PCR inhibitors (Moore and Dowhan 2007). The DNA was vacuum centrifuged to concentrate if necessary.

For all genets, at least 50 ng of high-quality genomic DNA was sent to Admera Health Biopharma for library preparation and sequencing. Paired end (2 X 150bp) sequencing libraries were prepared using the Kapa Hyper Prep minimal PCR kit and sequenced on the Illumina Novaseq X plus. A total of 27 samples, including 16 sexually produced offspring and 11 founders were sequenced targeting high coverage (∼50x) for inclusion in a broader, Caribbean-wide haplotype panel (Locatelli 2024) and for validation of shallow genotype calls (Table S2). A total of 157 samples comprising 153 sexually produced offspring (of which 16 were also sequenced at high coverage) and four founder genets, were sequenced targeting shallow coverage (∼3x, Table S3). Sequencing yielded an average of 67.5 million raw paired end reads per sample (range: 47.8 - 95.2 million) on average for the high coverage samples whereas the raw yield for the shallow coverage library was 6.7 million paired end reads per sample (range: 4.5 - 14.8 million) on average.

### Quality filtering, mapping and variant calling

All bioinformatic work was performed on the University of Southern California’s Center for Advanced Research Computing (CARC) system. First, sequencing adapters, low quality ends (<Q20), and 5’ bias were trimmed using Trim Galore v0.6.6 (Krueger 2015). Reads were then filtered to retain those with a 99% base call accuracy over at least 90% of the read using the fastx-toolkit v0.0.14 (Assaf and Hannon 2010). For the high coverage libraries, an average of 99.9% of reads remained after trimming (range: 99.7%-99.9%) and an average of 95.5% remained after quality filtering (range: 94.5%-96.3%). For the shallow libraries, an average of 99.7% of reads remained after trimming (range: 99.4%-99.9%) and an average of 89.6% remained after quality filtering (range: 85.7%-93.2%). Paired end reads where both mates passed trimming and filtering steps were distinguished from those where only one of the mates passed and both were independently mapped to a combined *A. palmata* (NCBI assembly: GCA_964030605.1) and Symbiodiniaceae genome reference (*Symbiodinium microadriaticum (Aranda et al. 2016)*, *Breviolum minutum* (Shoguchi et al. 2013), *Cladocopium goreaui* (Liu et al. 2018), *Durusdinium trenchii* (Dougan et al. 2024)) genome reference using the BWA-mem algorithm v0.7.17 (Li 2013) and then split into separate host and symbiont SAM files, retaining only host-specific files for further analyses. In total, mapped read percentages were high on average for both the high coverage (mean: 89%, range: 31-95%) and shallow library (mean: 90%, range: 15-95%) datasets, but six samples were removed from the shallow coverage dataset and two from the high coverage dataset due to low mapped read yields (< 2 million per shallow sample or < 50 million per high coverage sample) which corresponded to low coverage (<1x per shallow sample, ≤15x for high coverage sample) (all sexually produced offspring, Table S2,3). For the remaining 25 high quality, high coverage library samples, an average of 96.1 million reads mapped to the *A. palmata* genome (range: 55.8-144.9 million) yielding an average coverage of 33x (range: 18-52x). For the 151 remaining high quality shallow library samples, an average of 8.8 million reads mapped to the *A. palmata* genome (range: 2.8-13.9 million) yielding an average coverage of 3.5x (range: 1-5x). Paired and single-end mapped reads were then merged and PCR duplicates were marked using the Picard v2.26.2 MarkDuplicates tool (http://broadinstitute.github.io/picard). For both datasets, variant call files (VCF) were generated per sample using the multiallelic caller in bcftools v1.21 (Danecek et al. 2021) requiring a minimum map quality score of 40 and a minimum base quality score of 20 prior to merging.

### Genotype Imputation

A population-specific haplotype panel and linkage map for *Acropora palmata* (Locatelli 2024; Locatelli et al. 2024) was used to impute genotypes for the shallow coverage samples. One additional sample (AP66) was excluded from the sample set to be imputed due to its inclusion in the reference panel, leaving 150 shallow samples. Imputation was only performed for nuclear chromosomes (OZ034921.1-OZ034934.1). Prior to imputation, the shallow library VCF was further filtered to remove duplicate variants, invariant sites, and low allele count (AC < 3) variants, retaining only biallelic sites. A two-step imputation was performed following Hui et al. (Hui et al. 2020). First, Beagle v4.1 (Browning and Browning 2016) was used to estimate genotype probabilities (GP) from sample-specific genotype likelihoods referenced against the haplotype panel. Sites with a maximum genotype probability less than 99% (GP < 0.99) were set to missing. A second round of imputation was undertaken to impute missing sites using these high confidence genotypes in Beagle v5.4 assuming an effective population size of 10,000 individuals, as Ne values between 1-10,000 have been found to be nearly equivalent in terms of imputation error rate (Browning et al. 2018, 2021; Pook et al. 2020).

To assess performance of the genotype probability recalibration step and determine which post-imputation filtering threshold yielded the best accuracy, we compared genotype calls for replicate samples in the high coverage dataset with the raw, recalibrated and imputed genotypes in the shallow coverage dataset. Replicate samples included in the high coverage library (Table S4) were extracted from the high coverage VCF and shallow coverage post-imputation VCF files. The high coverage VCF sample-specific subset was further filtered to retain only biallelic sites genotyped in the shallow coverage post-imputation VCF files. The intersection of imputed sites that were also reliably genotyped in the high coverage dataset was used to assess accuracy by calculating the average Pearson’s correlation across sites and samples between the ‘true’ (i.e. high-coverage genotype calls at sites with ≥ 10x coverage) and the most probable imputed genotypes. We also calculated genotype concordance, or the proportion of identical genotype calls for both homozygotes and the heterozygote genotype across sites. We ultimately elected to retain only newly imputed sites with a dosage R^2^ greater than 99% (DR2 > 0.99) and sites with a genotype probability less than 99% (GP < 0.99) were again set to missing.

### Relatedness

Relatedness among batch cross samples was inferred using KING (Manichaikul et al. 2010) as implemented in plink2 (Chang et al. 2015). To allow for estimation of parental contributions, we generated a combined VCF dataset which in addition to the remaining shallow coverage samples (N=150) also included genotype calls for additional founders (N=11) and replicate sexually produced offspring (N=14) sequenced as part of the High Coverage dataset to verify relatedness thresholds using minimal filtering per best practice recommendations (https://www.kingrelatedness.com/manual.shtml). This merged dataset was filtered to remove low allele count (AC < 3) variants and retain only biallelic sites. The VCF was then converted to BED format using plink2 (Chang et al. 2015). Relatedness was estimated using the –make-king-table and –make-king square functions and the output was visualized using the dplyr, corrplot, and ggplot packages (Wickham 2016; Wickham et al. 2023) in R Studio v4.4.2 (R Core Team and Others 2023).

### Genetic Diversity

We calculated the average number of pairwise differences per site, or π, for Florida Keys founder genets (Upper and Lower Keys) and E-B Batch Cross offspring sequenced as part of the ‘high coverage’ dataset (Table S2) using pixy v2.0.0.beta8 (Korunes and Samuk 2021) with a sliding window size of 10 kb. A random subset of 5 individuals was selected for each ‘population’ (Upper Keys Founders, Lower Keys Founders and E-B Batch Cross), the maximum number available for the Lower Keys Founders, to minimize bias due to uneven sample size effects. Both variant and invariant sites were retained, but sites were filtered to retain at most biallelic records, with a per-sample depth greater than 10 reads but total depth less than 2.5x the mean coverage (<∼83x). An anova was used to test for differences in π among founder populations and the E-B Batch Cross offspring, followed by a Tukey’s HSD post-hoc test to investigate pairwise differences.

We also calculated the ID_risk_ statistic, which quantifies how long runs of homozygosity (ROH), reflective of the magnitude of inbreeding load, together with heterozygosity in non-ROH regions, reflecting the extent of recent inbreeding, can be used to predict a population’s risk of inbreeding depression (Kyriazis et al. 2025). As sample size does not impact estimates of ID_risk_, all remaining individuals were retained and sites were filtered to remove low allele count (AC < 3) variants and retain at most biallelic records, with a per-sample depth greater than 10 reads but total depth less than 2.5x the mean coverage (<∼83x). Length and ROH across individuals were detected using GARLIC v.1.1.6a (Szpiech et al. 2017), with an error rate of 0.001, and auto-detection of best window size and population allele frequencies estimated using exclusively adults.. Best practices for generating locations of centromeres for running GARLIC in non-model organisms were adapted from Szpiech et al. 2017. Further filtering and analyses of ROH were performed in R v. 4.5.2. Minimum Length of ROH (kb) was set 10 kb to exclude ancient population processes. To identify what length of ROH (kb) likely still harbors deleterious variation and is therefore functionally relevant in the existing population, runs of homozygosity were timed to their last common ancestor. Recombination rate was calculated using the maps generated from Locatelli et al. 2023 adapted to the *A. palmata* genome used in this study (NCBI assembly: GCA_964030605.1). The expected number of generations to a common ancestor (g) for an ROH of a given length (kb) was calculated using the following equation:

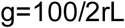

Where r is the recombination rate, and L is the length of ROH in kbp. ROH thresholds for capturing the functional effects of inbreeding were set using the assumption that recessive deleterious variants segregate for no more than 25 generations. IDrisk was calculated according to Kyriazis et al. 2025:

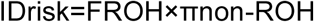

Where F_ROH_ is calculated as the fraction of the genome spanned by functionally-long ROH, and πnon-ROH is the heterozygosity in non-ROH regions. Heterozygosity in non-ROH regions of the genome was calculated using plink2 (--het, plink2/2.00a3.7LM, (Chang et al. 2015).

## Results

### Accuracy of imputed genotype calls derived from shallow whole-genome sequencing

We applied a two-step protocol developed for imputing genotypes from ultra-low coverage ancient DNA samples (Hui et al. 2020). The first step involves recalibrating raw genotype calls using genotype probabilities through comparison with known haplotypes, while the second step imputes genotypes at novel sites based on these higher confidence genotypes and the haplotype panel. To validate the accuracy of this approach, we sequenced a subset of individuals at both high (mean = 33x) and low (mean = 3.5x) coverage, then compared genotype accuracy and concordance. Accuracy of raw genotype calls in the low coverage sample set was low on average (Pearson’s r^2^ = 0.68) with heterozygote calls showing the lowest concordance across all alternate allele frequency bins (Fig. S1). Following genotype probability score recalibration, mean accuracy of genotype calls increased (Pearson’s r^2^ = 0.865) and heterozygote concordance improved (Fig. S2). Evaluation of accuracy across a range of imputation dosage R^2^ scores (DR2) revealed that the accuracy of novel heterozygote calls was improved with increasing DR2 and that accuracies were highest at the most stringent DR2 threshold (DR2>0.99, Fig. S3). Applying this threshold across all sites, we achieved a novel site imputation accuracy (Pearson’s r^2^) of 0.93 and a mean concordance of 0.95, 0.97 and 0.99 for homozygous reference, heterozygote, and homozygous alternate calls, respectively (Table S5). The average number of newly imputed sites per sample was 130,332 (range: 130,663-128,724, Table S3), representing 8% of the total 1,615,451 variable sites.

### Sexually produced restoration genets derived from batch crosses exhibit high relatedness

We calculated pairwise kinship coefficients to determine the degree of relatedness among offspring and putative founder parents. Twelve of the thirteen known technical duplicates (samples replicated in the high and low coverage sequence datasets) were accurately identified as clones based on elevated kinship scores (>0.354), with the thirteenth exhibiting a kinship score just below the prescribed threshold for clones (0.351, Table S6). An additional sixteen unexpected clonal relationships were also detected, all occurring among offspring from the E-B and E-H-SI Batch Crosses (Table S6). Of these, four clonal pairs were detected within E-B Batch Cross offspring, while two clonal pairs and two clonal triads were detected within E-H-SI Batch Cross offspring (Table S6), leading to genotypic richness values of 0.937 and 0.696 for the E-B and E-H-SI Batch Crosses, respectively. Finally, one additional clonal triad spanned the E-B and E-H-SI Batch Crosses, with AP72 from the E-B Batch Cross identified as a likely clone of both X-AP-12 and AP21 from the E-H-SI Batch Cross (Table S6).

Of the known parent-offspring relationships, 38/42 were accurately recovered based on kinship coefficients in the expected range of first-degree relationships (i.e. parent-child or sibling, 0.177-0.354, Table 1). Specifically, genet ML-LK10 (HG0543) from the high coverage dataset was a parent used in the LK-3 x LK-10 Biparental Cross. Thirty-eight Biparental cross samples were assigned kinship coefficients with ML-LK10 ranging from 0.185-0.279 (mean = 0.224, Table S7) in alignment with our expectation of a parent-offspring relationship. Of the remaining four unassigned offspring, two exhibit kinship coefficients with the parental genet just below the threshold for first degree relationships (0.17), one had a kinship coefficient reflective of a high-range second degree relationship (0.15) whereas the final, AP341, exhibited only third-degree relatedness (0.057, Table S7).

**Table 1.** Expected and observed parental relationships among founders and offspring from different batch crosses. MLG: Multi-locus genotype assignment from *Acropora* SNP-chip Galaxy Database obtained based on Parental IDs (Kitchen et al. 2020).

| Parental ID | Parental MLG | Parent Reef Origin | N offspring expected<sup>1</sup> | N offspring observed | Offspring Cross Origin (N) |
| --- | --- | --- | --- | --- | --- |
| ML-LK10 | HG0543 | Looe Key | 42 | 38 | LK-3 x LK-10 Biparental Cross |
| Apal-044 | HG0827 | Elbow Reef | >0 | 15 | E-B Batch Cross (14), 2014 Batch Cross (1) |
| Apal-074 | HG0822 | Elbow Reef | >0 | 47 | E-B Batch Cross (42), 2014 Batch Cross (2), 2015 Batch Cross (1), E-H-SI Batch Cross (2) |
| Apal-016 | HG1714 | Molasses Reef | 0 | 2 | 2015 Batch Cross (1), E-H-SI Batch Cross (1) |

<sup>1</sup>Our expectation for batch crosses is a parental contribution of greater than zero as records indicate gametes from parental contributors were not equalized at the time of fertilization and larvae from multiple batch crosses were subsequently pooled in 2017. If an equal number of gametes per parent had been mixed and there were no barriers to fertilization, the expected number of offspring per parent would be ~5.2 for the 2013 E-H-SI Batch Cross, ~0.6 for the 2014 Batch Cross and ~0.3 for the 2015 Batch Cross.

We next assigned parentage to unknown but expected relationships and explored the potential for unexpected parental and other relationships among batch cross offspring and wild adults. Two genets from the high coverage dataset, Apal-044 (formerly EL7, HG0827) and Apal-074 (formerly EL5, HG0822) were known to have contributed to batch crosses as parents from Elbow Reef (Table S1). Apal-044 was identified as a first-degree relative (∼parent) of 15 batch cross genets, predominantly originating from the 2017 batch crosses of parents from Elbow

Reef and Brewsters/Ball Buoy Reef (N=14) while Apal-074 was identified as a first-degree relative of 42 offspring from the E-B Batch Cross pool, in addition to first-degree relationships with offspring from batch crosses in other years again known to involve Elbow Reef parents (Table 1, S8, S9). When examining the intersection of these two parent-offspring lists, 14/15 of the Apal-044 offspring are contained within the Apal-074 list and an independent assessment of their pairwise Kinship coefficients (range: 0.183-0.32) establishes they are relatives in the first degree, confirming their assignment as full siblings (Table S10). We also detected two unexpected first degree relationships between founder genet Apal-016 from Molasses Reef and offspring from two independent batch crosses including Elbow Reef parents (Table 1, S11), a distance of nearly 19 km. Interestingly, a first-degree kinship on the order of a full sibling relationship was also detected between AP-074 and two other wild adults in Mote’s holdings, ML-SK-4 and ML-WDR-2 (Table S8) and as a result, similar first-degree kinship relationships were detected between these two founders and offspring of AP-074 (Table S12, S13). Whereas strong negative relationships, which may indicate population structure between individuals, were detected between one E-B Batch Cross genet, AP55, and all other E-B Batch Cross offspring (Table S14).

Examination of the full distribution of kinship coefficients within batch crosses indicates a high degree of relatedness overall and a substantial proportion of full-sibling relationships within each, despite the multiparent nature of the E-B and E-H-SI Batch Crosses (Fig. 2, Fig. S4). Hierarchical clustering analysis of pairwise kinship coefficients identified five major groups (Fig. S5) which reflect sets of full-siblings based on prescribed thresholds for first-degree (i.e. full sibling, 0.177-0.345), second-degree (i.e. half-sibling, 0.0884-0.177) and third-degree (i.e. first cousin, 0.0442-0.0884) relationships (Manichaikul et al. 2010). The E-B Batch Cross, a pool of three different multi-parent larval fertilizations (Table S1), is largely made up of just three full-sib groups, whereas the E-H-SI Batch Cross is composed almost entirely of full-siblings, despite there being a maximum of 10 possible unique full-sibling groups among the five parents used in the cross (Fig. 2). Moreover, relatedness remains high among regionally distinct batch crosses, as evidenced by offspring from the Lower Keys Biparental Cross (LK3 x LK10) exhibiting kinship coefficients with both Upper Keys Batch Crosses on the order of half-sibling to first cousin relationships (∼2^nd^ and 3^rd^ degree relatives, Fig. 2).

**Figure 2.**
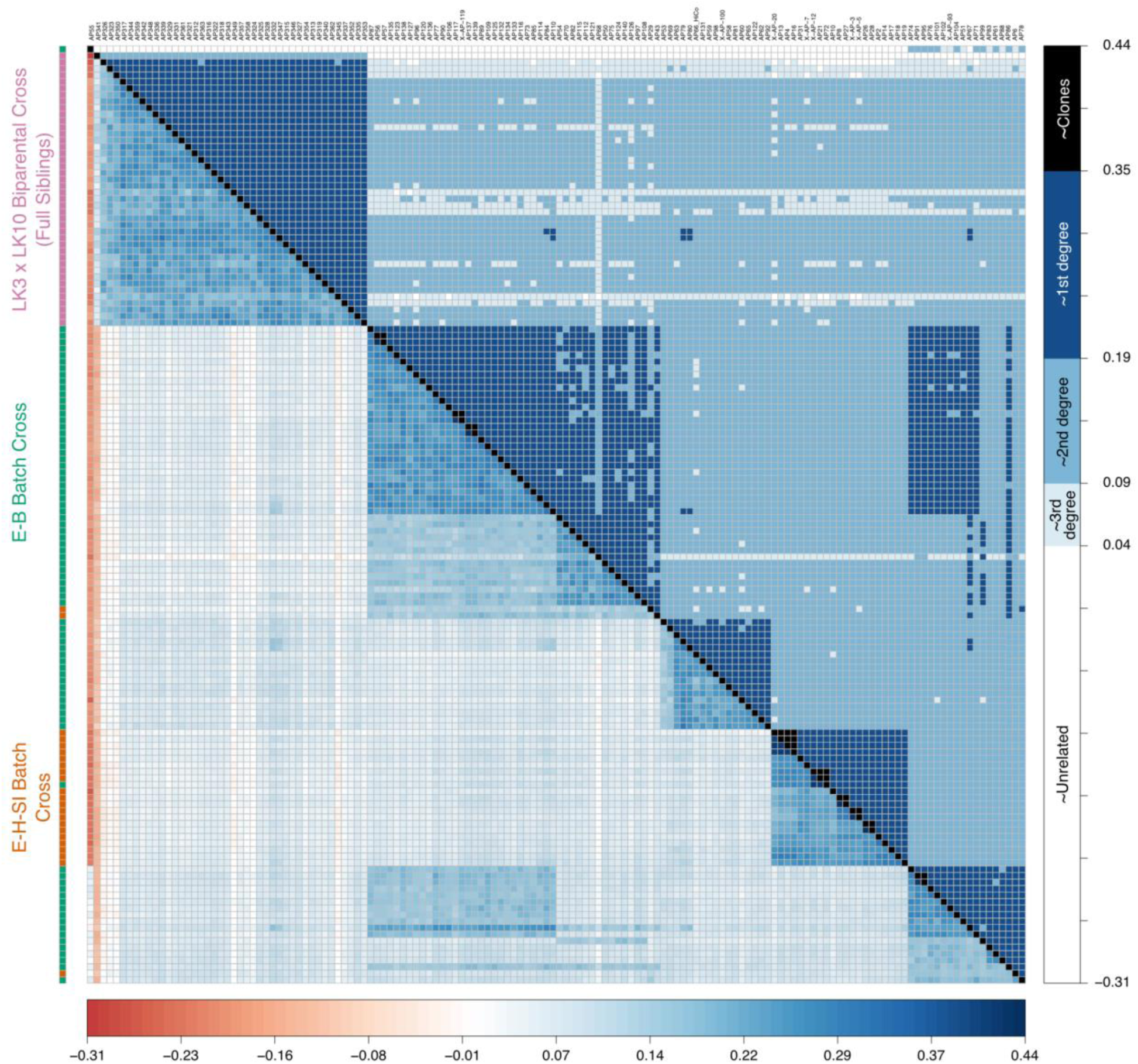
Matrix of pairwise kinship coefficients among batch cross offspring colored based on their absolute value (below diagonal) or binned by approximate relatedness thresholds (above diagonal). Samples were clustered using Ward’s method to limit within-cluster variance and order is identical in rows and columns. Rows are labeled by color according to their cross of origin.

### Minor reduction in genetic diversity, but low risk of inbreeding depression

We evaluated genome-wide nucleotide diversity using a subset of samples sequenced at high coverage to determine whether sexually produced recruits from the E-B Batch Cross differed from founder genets. Estimates were similar in magnitude, but significantly different (F_2,90564_ = 5.096, P=0.006, Fig. 3A). E-B Batch Cross genets had the lowest levels of diversity on average (mean π ≈ 0.00285, range: 0-0.135) followed by Upper Keys Founders (mean π ≈ 0.00293, range:0-0.110) and Lower Keys Founders (mean π ≈ 0.00299, range:0-0.130, Tukey’s HSD = 0.13). Pairwise comparisons revealed a significant difference in π between E-B Batch

**Figure 3.**
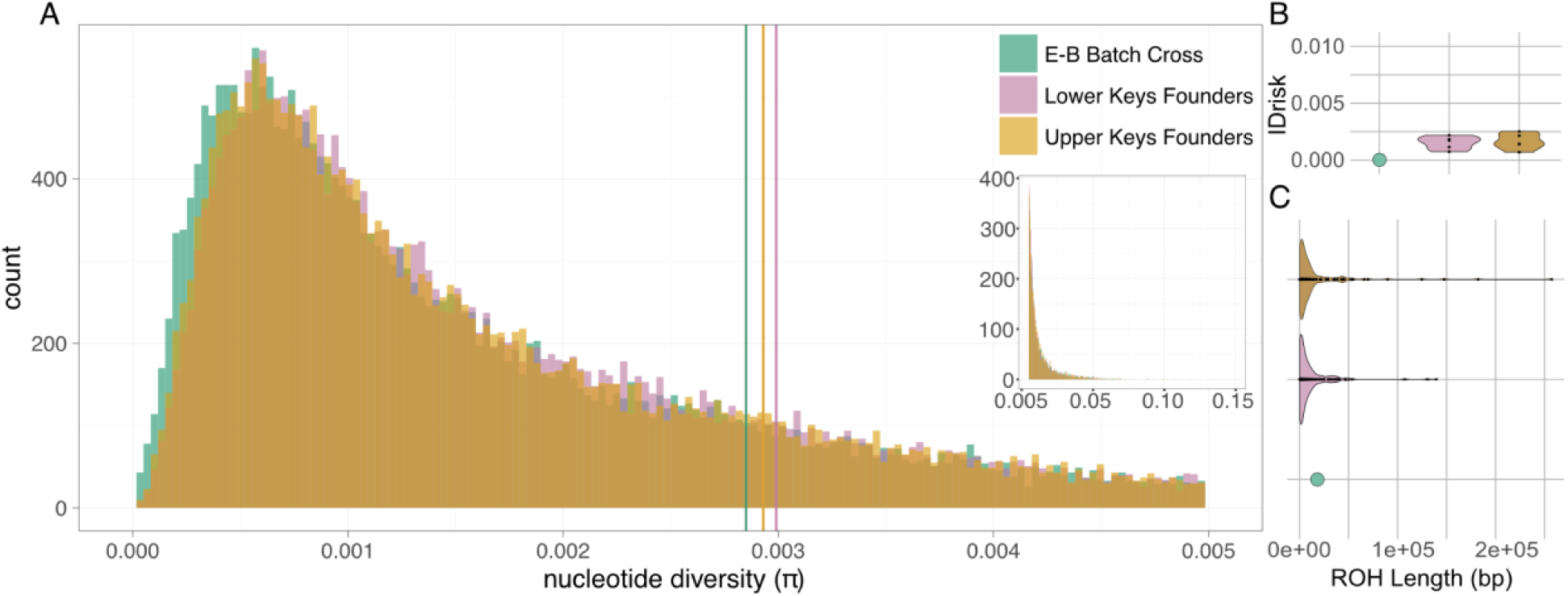
(A) Histogram of pairwise nucleotide differences (π) estimated for a random subset of five E-B Batch cross genets (EB), the five Lower Keys founders (LK) and five Upper Keys (UK) founders sequenced at high coverage. Vertical lines indicate population means. Note that distribution in main panel is truncated to π < 0.005 and the distribution for values up to 0.15 is shown in the inset. (B) Violin plots of inbreeding depression risk (IDrisk) for E-B Batch Cross and founder genets by population. Note that E-B Batch Cross is represented by only a single individual and point size has been artificially increased for visual clarity. Values less than 0.05 are defined as ‘low risk’ by Kyrazis et al. 2025. (C) Violin plots showing length distribution of runs of homozygosity (ROH) for E-B Batch Cross and founder genets by population. Note that E-B Batch Cross is represented by only a single individual and point size has been artificially increased for visual clarity.

Cross genets and Lower Keys Founders (Tukey’s HSD P_adj_ = 0.004) but not between E-B Batch Cross genets and Upper Keys Founders (Tukey’s HSD P_adj_ = 0.13) or between founder groups (Tukey’s HSD P_adj_ = 0.43).

Inbreeding depression risk (IDrisk), calculated as the fraction of the genome spanned by functionally-long runs of homozygosity (ROH) multiplied by heterozygosity in non-ROH regions was low (defined as values <0.05, (Kyriazis et al. 2025), less than 0.0025 on average for all samples (Fig. 3B), and almost unquantifiable in E-B Batch Cross genets due to undetectable ROH in all but one genet (Fig. 3C).

## Discussion

Scaling up of larval-based coral restoration methods, in which offspring of natural mass spawning events are either reared in aquaria and incorporated into nursery collections (Koch et al. 2022) or directly recruited onto natural reefs (Harrison et al. 2021), is a major priority for reef restoration efforts globally (Hardisty et al. 2019; Randall et al. 2020; Vardi et al. 2021; Banaszak et al. 2023). Proposed advantages of this approach include harnessing the r-selected life-history of coral, which naturally produce high numbers of larvae, to maximize the number of surviving recruits in support of demographic and ecosystem recovery (Hardisty et al. 2019; Harrison et al. 2021). In addition, recombination generates new allelic combinations in offspring, creating new genetic variation upon which selection can act (Vardi et al. 2021), which again may be magnified by the high fecundity of coral (Williams 1975). However, there are also risks including known gamete incompatibilities among potential parents (Baums et al. 2013; Miller et al. 2018; Koch et al. 2022) and bottlenecks during the larval culturing and recruit grow-out phases which can reduce genetic diversity in the final offspring pool (Baums et al. 2022; López-Nandam et al. 2022).

Here we show that while a large number of offspring genets can be successfully reared from batch crosses for use in reef restoration, these individuals exhibit very high relatedness, meaning that the realized contribution of each parent is lower than the apparent number of individuals included in the cross. For example, although 80 individual offspring were propagated from the E-B Batch Cross larval pool, we found they comprise only three full-sibling groups, despite being derived from three independent batch fertilizations using gametes from 3-5 different parents, which combinatorially could have produced a maximum of 6-10 full-sibling groups each. Captive breeding programs for terrestrial species have long recognized the importance of managing genetic diversity, leading to routine use of pedigrees and other molecular markers to track parentage and limit inbreeding (Lacy 1994; Ivy and Lacy 2010).

Although we detected reduced genetic diversity in E-B Batch Cross offspring relative to some parental founder populations, the difference is not yet large in absolute terms, a pattern further reinforced by a low inbreeding depression risk. Therefore, there is still time to minimize the potential for future inbreeding in restored coral populations. Below we consider possible explanations for the elevated relatedness observed in these captive populations of sexual recruits and ways practitioners can mitigate genetic risk through careful restoration planning.

### Mechanisms contributing to elevated relatedness of sexual recruits

There are multiple mechanisms which can lead to unequal parentage within batch cross offspring, resulting in the predominance of specific parents as observed here (Fig 2, Table 1). Fertilization can occur rapidly (Nozawa et al. 2015; Chamberland et al. 2025), and if gamete additions to batch crosses occur sequentially, priority effects may lead to over-representation of specific offspring pairs. In addition, fertilization success differs for different pairwise crosses of *A. palmata*, suggesting the existence of intrinsic genetic incompatibilities even among unrelated founder individuals (Baums et al. 2013, 2022; Miller et al. 2018). Through parentage analysis of four-parent batch crosses, Baums et al. (2013) report over-representation of specific biparental combinations as early as 1-2 days post-fertilization. This could be reflective of both pre- and post-zygotic barriers to reproduction. Sperm competition and variation in gamete affinities has been repeatedly observed in mass spawning marine invertebrates, which is likely attributable to genetic variation in gamete recognition proteins among individuals (Palumbi 1999; Levitan and Ferrell 2006; Levitan 2012, 2018). This would preclude successful fertilization in the first place, a phenomenon which has been observed in *A. palmata*’s sister species, *A. cervicornis* (Koch et al. 2022), and which may be one explanation for the large range of fertilization success observed in specific biparental crosses (Baums et al. 2013; Miller et al. 2018). Incompatibilities can also arise during subsequent larval development. Larval mortality rates are high for many marine invertebrates (Rumrill 1990); leading to high variability or sweepstakes reproductive success in natural populations (Hedgecock and Pudovkin 2011). Whether the result of priority effects arising from time-lagged additions of gametes to the batch cross, intrinsic genetic incompatibilities, or laboratory induced selection, the outcome remains the same: certain genetic combinations are lost from the larval population leading to reduced genetic diversity in offspring cohorts.

While clonal reproduction is a well-established life-history strategy in adult coral populations (Manzello et al. 2019; Drury et al. 2019), it is also possible for developing zygotes to be broken up by shear forces in the early stages of cell division, resulting in clonal reproduction during the larval life stage (Heyward and Negri 2012). In addition, technical biases such as low coverage, in combination with the low overall genetic diversity present in the species, can also lead to inflated kinship values, wherein siblings can be erroneously classified as clones. In the present crosses, we observed elevated clonality in the batch crosses, particularly in the E-H-SI cross which exhibited genotypic richness of only 0.696, which while not as low as some founder populations in Florida, is within the range of observed values for clonally reproducing adult populations (Baums et al. 2006). Alternatively, or in addition to this explanation, accidental mislabeling during intentional asexual propagation of offspring genets could also lead to apparent clones (Lirman et al. 2010; Koch et al. 2022). For example, we observed a first-degree relationship (i.e. parent-offspring or full-sibling) between an E-B Batch Cross genet (the product of Upper Keys Founder genets) and two distinct founder genets from the Lower Florida Keys, populations separated by 185-km. It is biologically implausible that sperm could traverse this distance within the prime fertilization window, as fertilization success is known to decline rapidly over the scale of meters in natural populations of acroporids (Mumby et al. 2024; Ricardo et al. 2026). Similarly, two nominally E-H-SI batch cross genets have a parentage assignment to AP-074, which according to records was not a contributor to the 2013 E-H-SI cross and cluster as full sibs with E-B batch cross genets to which AP-074 did contribute gametes. A more likely explanation is that ramets were simply mislabeled at some point during the *ex situ* propagation process. Tracking hundreds to thousands of unique coral genets, each represented by potentially thousands of ramets within a production nursery, is an extreme logistical challenge facing coral restoration practitioners. This highlights the need for optimizing and standardizing processes for inventory tracking in restoration in support of genetic management objectives.

### Low risk of inbreeding depression at present

Our genome-wide estimate of genomic diversity of 0.285% is lower, but on par with that reported for an Indo-Pacific congener, *A. millepora*, (0.363%, (Prada et al. 2016; Cramer et al. 2020; Fuller et al. 2020; Cooke et al. 2020). If we assume a similar mutation rate of 4 x 10^-9^ per base per generation (Matz et al. 2018), this corresponds to a historical effective population size (Ne) of around 1.78 × 10^5^, although caution is warranted since this estimate is highly dependent on mutation rate and is an inaccurate estimate of contemporary Ne. A recent regional estimate for the sister species, *A. cervicornis*, puts contemporary Ne on the order of ∼70 individuals (Duffin et al, in prep.). Taken together, this means that existing founders represent a historically diverse gene pool, which has experienced a comparatively recent demographic collapse (Cramer et al. 2020). While large population sizes increase the genetic risk arising from inbreeding following population contraction (Charlesworth and Willis 2009), and even further population contraction was recently documented in Florida (Williams et al. 2020; Manzello et al. 2025), we see no evidence of inbreeding in founder genets, or in this F1 cohort of sexually produced offspring. Indeed, runs of homozygosity were nearly undetectable in the E-B Batch Cross representatives, likely due to recombination during meiosis. However, our sampling was conducted prior to the 2023 mortality event, and the high relatedness in these F1 restoration cohorts combined with the recent functional extinction of founders in Florida underscores the importance of minimizing inbreeding risk in future breeding and outplanting designs through careful genetic management.

### Recommendations for maximizing genetic diversity of sexually produced coral populations

Maximizing genetic diversity of coral outplants is a major goal of reef restoration programs (Baums et al. 2019, 2022). For programs employing sexual reproduction, minimizing kinship or pairwise relatedness among sexually produced offspring, has been recommended as one approach to maximizing genetic diversity of restoration populations (Baums et al. 2022), although questions remain as to optimal outplant designs which both incorporate novel allelic combinations resulting from sexual reproduction while also minimizing inbreeding risk. In line with prior observations of unequal parental contributions in multi-parent batch crosses (Baums et al. 2013; López-Nandam et al. 2022), we show that sexually produced restoration lineages derived from wild founders can suffer from the same biases, although this bias may be species specific (Dallmeyer-Drennen et al. 2026) and can be mitigated through process adjustments such as simultaneous mixing of gametes rather than sequential addition to batch crosses. While there is an argument to be made for the demographic importance of these genets regardless of their relatedness given that *A. palmata* is now functionally extinct on Florida’s reefs (Manzello et al. 2025), there are also risks to outplanting highly genetically related cohorts in the absence of natural founders.

A major goal of reef restoration is to establish self-sustaining reefs, meaning achieving natural sexual reproduction among outplanted genets is a priority (Baums et al. 2019). However, if a cohort of Batch Cross genets were outplanted in proximity, subsequent sexual reproduction would yield a high probability of offspring resulting from full sibling matings. While these hypothetical inbred offspring would then hopefully disperse to other reefs, if the same parental founder or Batch Cross genets are repeatedly used and shared among regional practitioners, there is also a possibility that these hypothetical inbred offspring could recruit to reefs restored with other full or half sibling genets or even ramets of their founder parents, leading to additional inbreeding risk in future generations. While this scenario is unlikely for several reasons, not least of which is the pervasive absence of successful larval recruitment in Caribbean acroporids (Williams et al. 2008), we believe that this scenario can nevertheless be avoided without significantly negatively affecting outplanting yield or effort through integrated genetic management and larval propagation protocols.

Substantial progress has been made in developing priorities and best-practice recommendations for quantifying genetic diversity, rebalancing abundances, and distributing ramets of founder genets among nursery holdings based on a recent population status assessment for *Acropora palmata* (Rodriguez-Clark et al. 2025) and we re-emphasize the need to adhere to these recommendations. Minimizing overall relatedness, by tracking parentage and prioritizing under-represented founders in crossing designs, is paramount. Ensuring that new breeders are regularly rotated into sexual production pipelines and retiring previous parents, such as the Elbow Reef genets (Table 1), can be done even in the absence of specific pedigrees (Rodriguez-Clark et al. 2025). When funding is available to support genotyping, Baums et al. (Baums et al. 2022) have outlined multiple methods that can be applied, and here we show that shallow whole genome sequencing is a cost-effective approach. Outplanting designs should take relatedness into account to minimize the risk of future inbreeding by placing close relatives at a sufficient distance from one another to prevent cross-fertilization (Baums et al. 2022; Rodriguez-Clark et al. 2025). Given that restoration is also occurring over a large spatial scale, possible dispersal kernels for source populations should also be considered to minimize inadvertent recruitment of close relatives to downstream populations, assuming that the current larval recruitment crisis can be resolved.

## Conclusions

Given the high relatedness observed in restoration lines of sexually produced coral both within and among cohorts, we advocate for coordinated genetic management planning among independent restoration organizations to limit the risk of inbreeding in future restoration outplanting populations. At a minimum, there should be regional coordination around rotation of broodstock to ensure annual offspring pools are unrelated in space and time. If offspring are to be incorporated into asexual propagation programs, we recommend regular genotyping of ramets and propagating only the most unrelated individuals for outplanting. By validating a genotype imputation approach, we show that accurate genome-wide genotyping can be achieved using more cost-effective shallow coverage (3-5x) whole genome sequencing. In addition, optimization and standardization of methods for tracking genet inventory can help limit unintentional overpropagation of clonal lineages. Taken together, careful genetic management planning and coordination among restoration partners can maximize the benefits and minimize the risks of sexual propagation approaches to reef restoration (Vardi et al. 2021).

## Data Accessibility

All raw sequence data have been archived on NCBI’s SRA under PRJNA1078365. Scripts necessary to replicate bioinformatic processing steps and subsequent statistical analyses and data visualizations can be found at https://github.com/ckenkel/ApalConGen (Zenodo DOI to be generated).

## Acknowledgements

Work was conducted under permits to Mote: FKNMS-2021-172, SAL-21-1724-SCRP and SAL-22-2406-SCRP and CRF: FKNMS-2019-012, SAL-23-2525-SCRP. We thank Phanor Montoya Maya for managing the CRF nursery collection. We also thank Eleftherios Karabelas for contributing to the execution of the project and data collection efforts. CDK, IBB, RSW, MWM and EMM are members of the Coral Restoration Consortium’s Genetics Working Group while CDK, MWM and EMM serve on the US Acropora Recovery Implementation Team, which is a formal advisory body to the US National Marine Fisheries Service. Funding for this study was provided by a NOAA Ruth D. Gates Coral Restoration Innovation Grant #NA21NMF4820300 to CDK and EMM.

## Supplementary Material

**Table S1.** Spawning records for coral which likely contributed to the parentage of each cohort. Note that in 2017 three separate fertilizations were recorded, two on 10 August composed of gametes from Elbow reef founders (Batch 1) and gametes from Biscayne National Park founders (Batch 2) and one on August 11 composed of gametes from three Elbow reef genets. All larvae were subsequently pooled at Mote as the 2017 cohort.

| Genet Name | Genet ID (SNP chip) | Cross Year / Day / Fertilization Batch |  |  |  |  |  |  |
| --- | --- | --- | --- | --- | --- | --- | --- | --- |
|  |  | 2013 | 2014 | 2015 | 2017 |  |  | 2020 |
|  |  |  |  |  | 10 Aug |  | 11 Aug |  |
|  |  |  |  |  | Batch 1 | Batch 2 |  |  |
| EL Green | HG0083 |  | X |  | X |  | X |  |
| EL Yellow | HG0793 | X | X | X | X |  |  |  |
| EL Orange | HG0822 |  | X | X | X |  |  |  |
| EL White | HG0827 | X | X |  | X |  |  |  |
| EL Blue | HG1548 or HG0822 |  | X | X | X |  | X |  |
| EL Unk | HG0796 or HG0787 |  |  |  |  |  | X |  |
| HS | HG0004 | X |  |  |  |  |  |  |
| SI Orange | HG0784 | X |  |  |  |  |  |  |
| SI Blue | HG0197 | X |  |  |  |  |  |  |
| BNP Ball 1 | NA |  |  |  |  | X |  |  |
| BNP Ball 2 | NA |  |  |  |  | X |  |  |
| BNP Brewster | NA |  |  |  |  | X |  |  |
| AP LK-3 | HG0186 |  |  |  |  |  |  | X |
| AP LK-10 | HG0543 |  |  |  |  |  |  | X |

**Table S4.** List of samples used for imputation validation and their corresponding genet identifiers.

| High Coverage SampleID | Shallow SampleID | Genet ID (SNP chip) | Genet Name |
| --- | --- | --- | --- |
| 21073D-02-25_S164_L003 | 21073D-03-38_S70_L003 | HG1093 | AP54 |
| 21073D-02-26_S110_L003 | 21073D-03-39_S71_L003 | HG1118 | AP55 |
| 21073D-02-27_S112_L003 | 21073D-03-40_S72_L003 | HG1045 | AP56 |
| 21073D-02-28_S113_L003 | 21073D-03-47_S79_L003 | HG1129 | AP63 |
| 21073D-02-31_S120_L003 | 21073D-03-59_S88_L003 | HG1077 | AP75 |
| 21073D-02-32_S108_L002 | 21073D-03-60_S89_L003 | HG1106 | AP76 |
| 21073D-02-34_S0_L001 | 21073D-03-70_S98_L003 | HG1090 | AP86 |
| 21073D-02-35_S103_L002 | 21073D-03-73_S101_L003 | HG1096 | AP89 |
| 21073D-02-36_S114_L003 | 21073D-03-75_S103_L003 | HG1068 | AP91 |
| 21073D-02-37_S0_L001 | 21073D-03-27_S60_L003 | HG0600 | AP30 |
| 21073D-02-39_S93_L002 | 21073D-03-12_S80_L005 | missing | AP14 |
| 21073D-02-40_S96_L002 | 21073D-03-07_S48_L003 | missing | AP8 |
| 21073D-02-30_S166_L003 | 21073D-03-128_S142_L003 | missing | AP327 |

**Figure S1.**
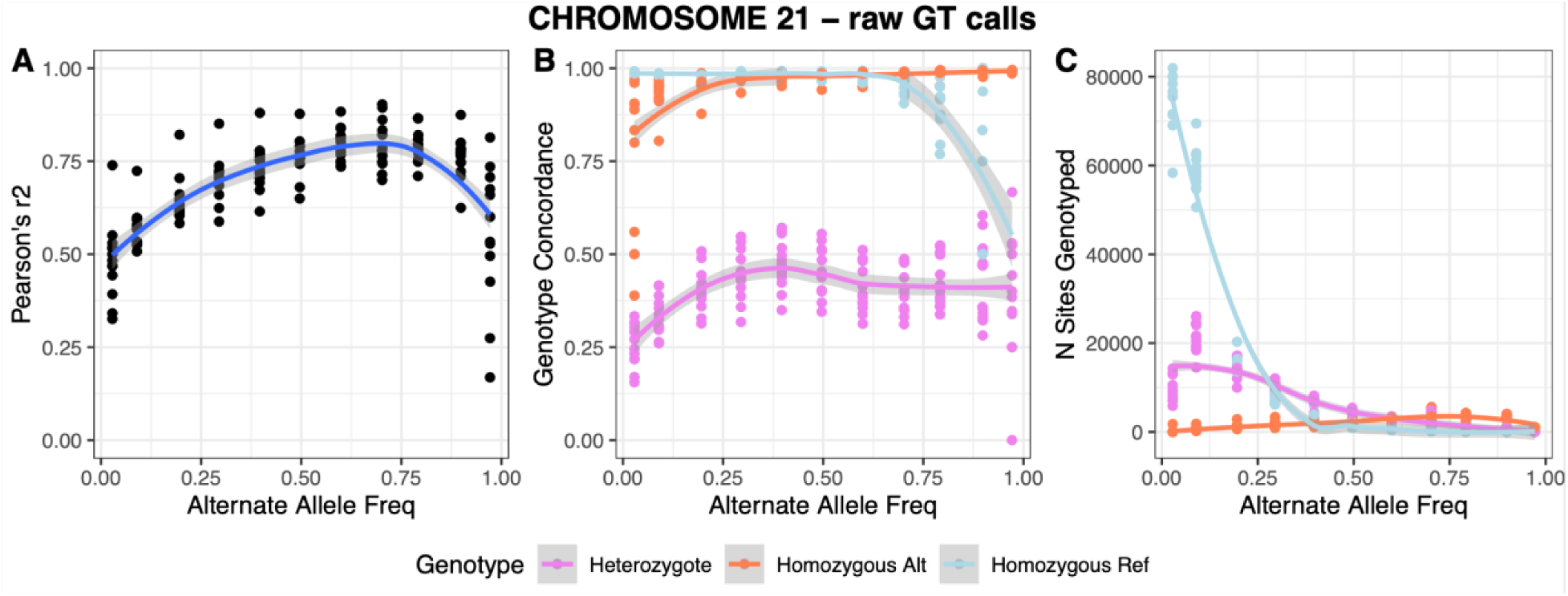
Accuracy of raw genotype calls (A), genotype concordance (B) and the number of sites genotyped (C) as a function of alternate allele frequency (AF) for Chromosome 21 relative to high-confidence calls (read depth > 10) in the high coverage dataset.

**Figure S2.**
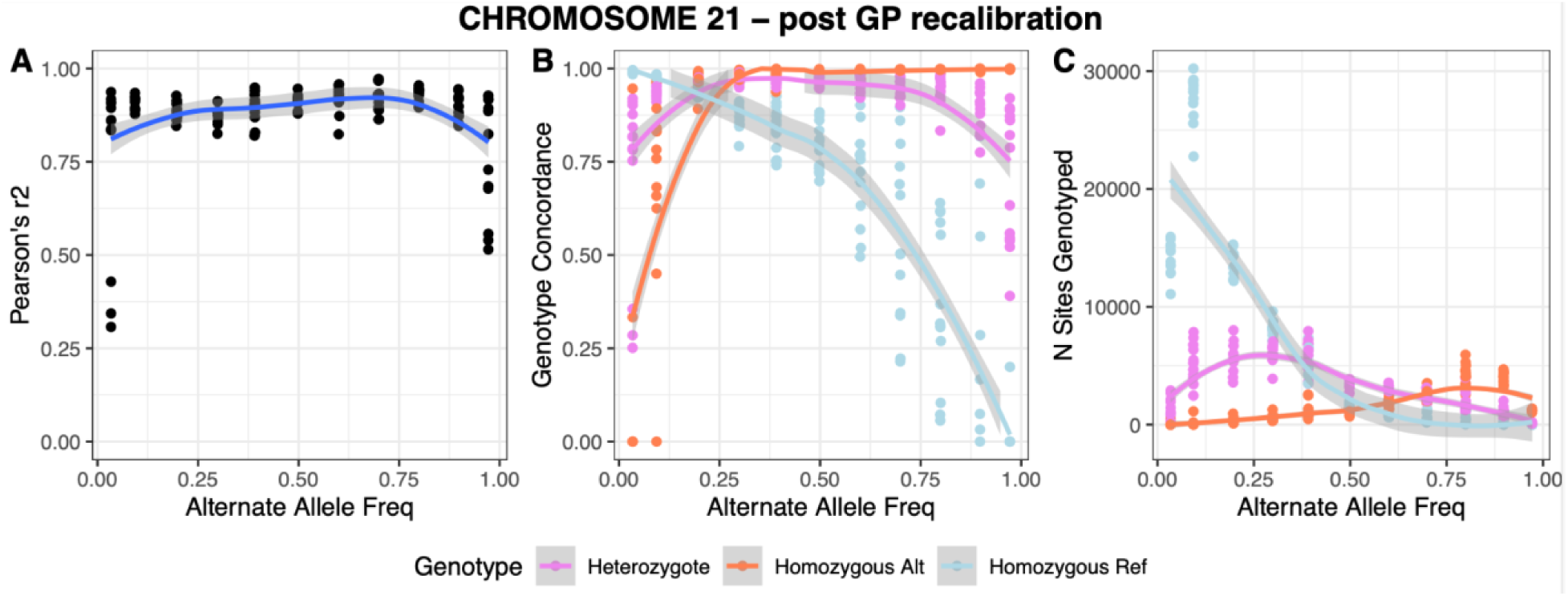
Accuracy of genotype-probability recalibrated genotype calls (A), genotype concordance (B) and the number of sites genotyped (C) as a function of alternate allele frequency (AF) for Chromosome 21 relative to high-confidence calls (read depth > 10) in the high coverage dataset.

**Figure S3.**
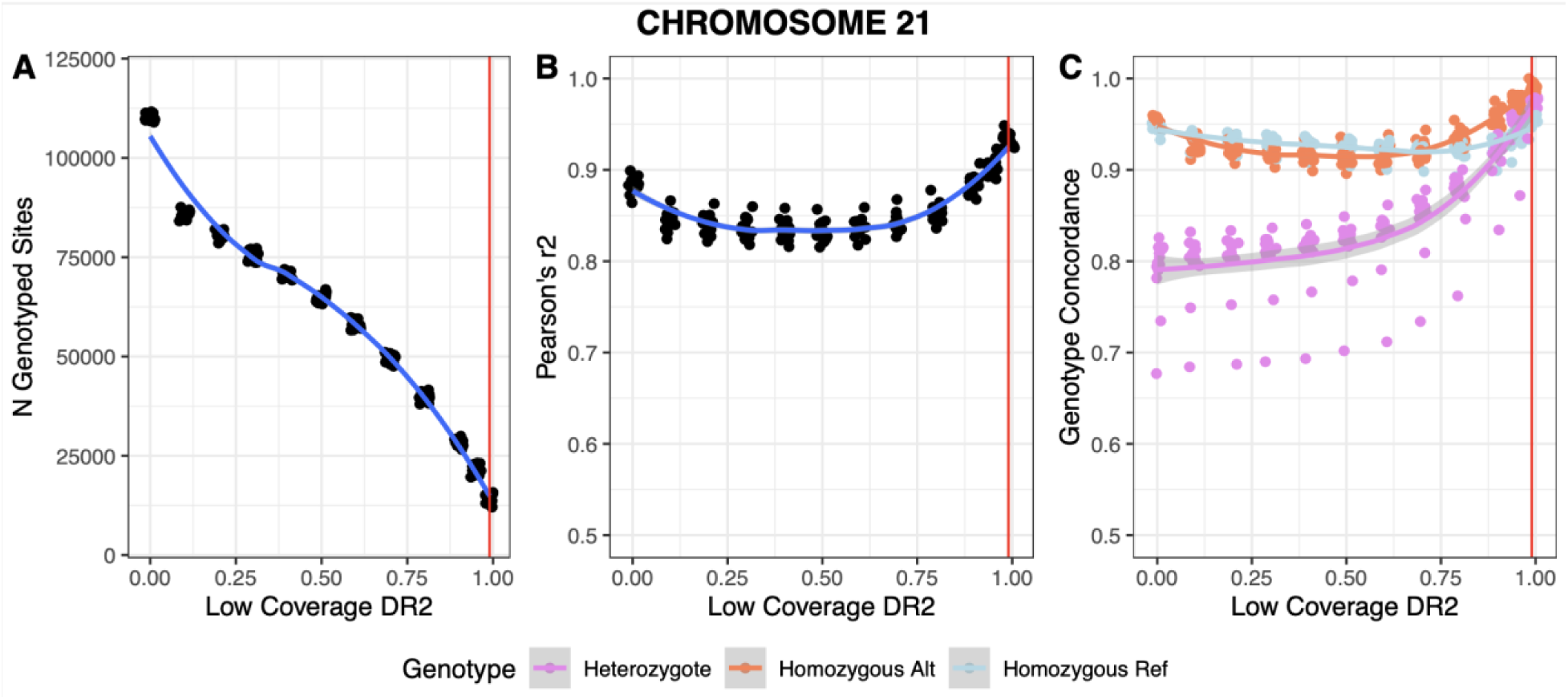
The number of imputed sites genotyped (A), imputation accuracy (B) and genotype concordance (C) as a function of imputation DR2 score for Chromosome 21 relative to high-confidence calls (read depth > 10) in the high coverage dataset. The red vertical line indicates the chosen DR2 filtering threshold of 0.99.

**Table S5.** Imputation accuracy summary statistics by chromosome.

| Chromosome | Correlation between high coverage calls and raw low coverage genotypes (Pearson's $r^2$) | Correlation between high coverage calls and GP-recalibrated genotypes (Pearson's $r^2$) | Correlation between high coverage calls and novel imputed genotypes (Pearson's $r^2$) | Imputation Concordance for sites with DR2 ≥ 0.99<br>HomRef : Het : HomAlt | N Imputed Sites |
| --- | --- | --- | --- | --- | --- |
| OZ034921.1 | 0.686 | 0.878 | 0.93 | 0.95 : 0.97 : 0.99 | 13,927 |
| OZ034922.1 | 0.702 | 0.878 | 0.94 | 0.96 : 0.97 : 0.99 | 9,804 |
| OZ034923.1 | 0.666 | 0.871 | 0.93 | 0.95 : 0.98 : 0.99 | 10,669 |
| OZ034924.1 | 0.698 | 0.883 | 0.95 | 0.97 : 0.97 : 0.99 | 9,475 |
| OZ034925.1 | 0.669 | 0.834 | 0.91 | 0.95 : 0.96 : 0.99 | 9,340 |
| OZ034926.1 | 0.673 | 0.854 | 0.92 | 0.95 : 0.97 : 0.99 | 10,414 |
| OZ034927.1 | 0.671 | 0.862 | 0.92 | 0.96 : 0.95 : 0.98 | 10,879 |
| OZ034928.1 | 0.711 | 0.878 | 0.94 | 0.96 : 0.95 : 0.99 | 7,183 |
| OZ034929.1 | 0.681 | 0.876 | 0.94 | 0.96 : 0.98 : 0.99 | 9,770 |
| OZ034930.1 | 0.682 | 0.859 | 0.94 | 0.96 : 0.98 : 0.99 | 7,903 |
| OZ034931.1 | 0.691 | 0.849 | 0.92 | 0.95 : 0.97 : 0.99 | 6,534 |
| OZ034932.1 | 0.674 | 0.850 | 0.91 | 0.95 : 0.97 : 0.99 | 8,152 |
| OZ034933.1 | 0.660 | 0.865 | 0.95 | 0.96 : 0.97 : 0.99 | 8,388 |
| OZ034934.1 | 0.664 | 0.879 | 0.94 | 0.96 : 0.97 : 0.98 | 8,627 |

**Figure S4.**
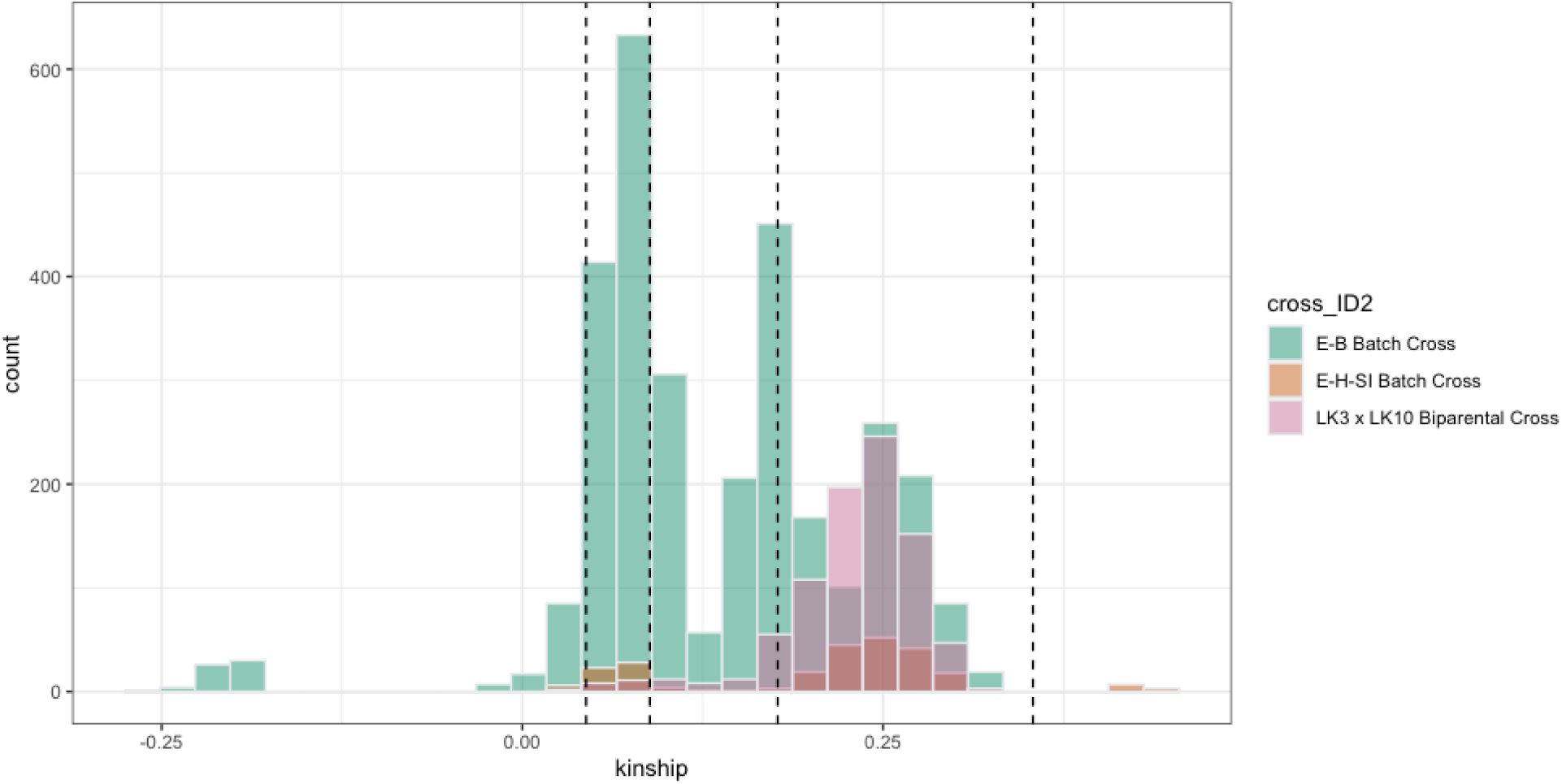
Histogram of kinship coefficients for within-batch cross comparisons. Vertical lines demarcate the prescribed coefficient ranges for identifying duplicates/twins (>0.354), first-degree (e.g. parent-offspring or full siblings, 0.177-0.354), second-degree (e.g. half-siblings, aunts/uncles, 0.0884-0.177), third-degree relationships (e.g. first cousins, 0.0442-0.0884) <u>(Manichaikul et al. 2010)</u>.

**Figure S5.**
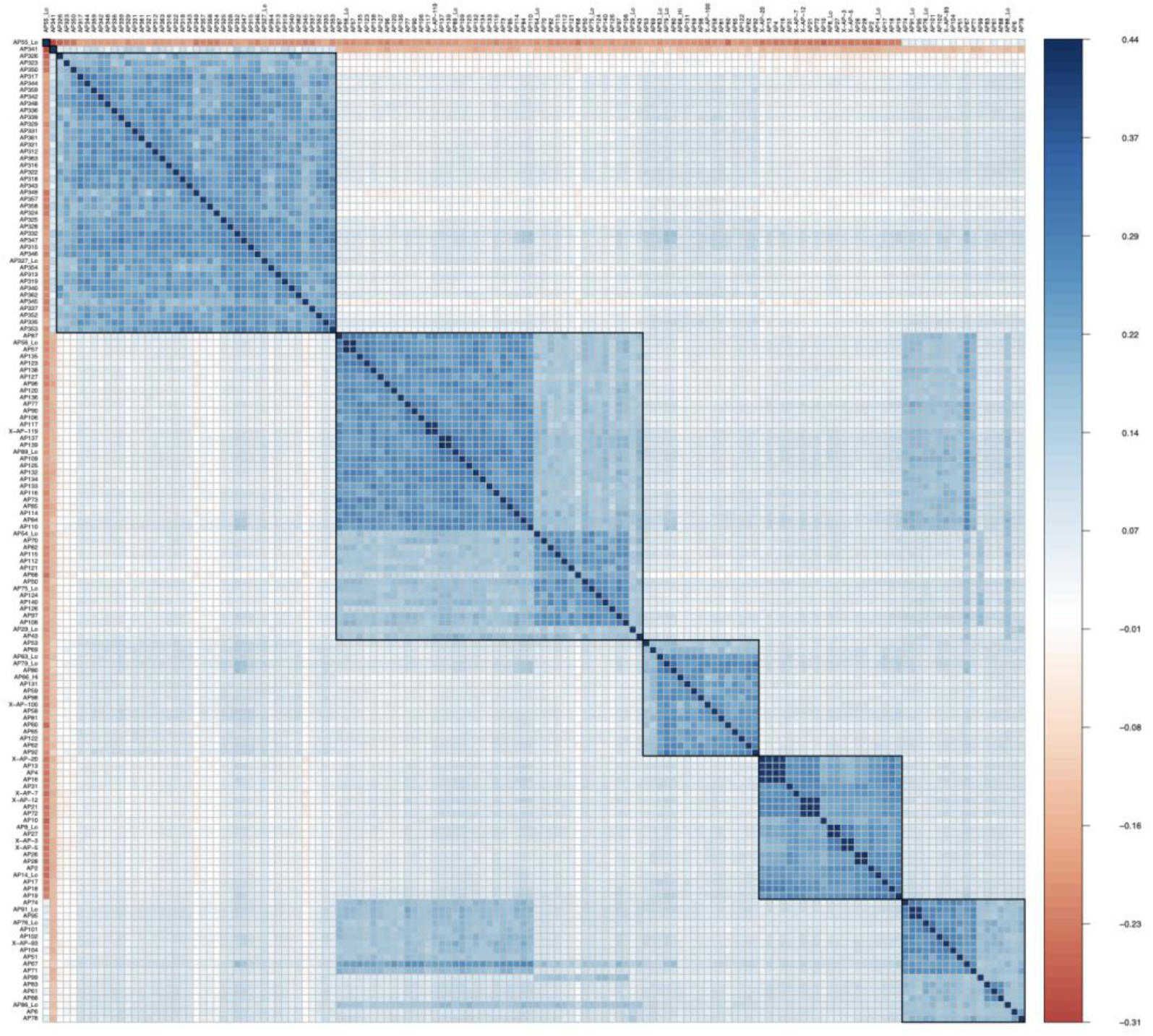
Correlation plot of the hierarchically clustered pairwise kinship coefficients between samples using Ward’s method to limit within-cluster variance.

